# Screening bacterial isolates for phosphate Solubilizing capability in a ferruginous ultisol in Benin City, Edo State Nigeria

**DOI:** 10.1101/2020.09.30.320952

**Authors:** S.I. Musa, Beckley Ikhajiagbe

## Abstract

Phosphorus is a major growth-limiting nutrient which plays important biochemical role in photosynthesis, respiration and several other processes in the living plant. It is widely distributed in minerals as phosphates. It reacts easily with Fe^3+^ in ferruginous ultisols and therefore not bioavailable for plant usage. Many bacteria have the ability to solubilize phosphate minerals and make it bioavailable to plants. Thus this research investigates the culturable bacterial composition of ferruginous ultisol, comparative to control soils as well as the phosphate solubilizing capabilities of the isolates for future use in soil improvements. Six soil samples of different ferruginous levels and a control were assayed for physicochemical parameters prior to the experiment. Culturable bacteria as well as the phosphate solubilizing bacteria (PSB) were assayed in Pikovskaya’s medium at 27°C with 7.5 pH for 7days. Six distinct isolates were observed which proved to be *Proteus spp*., *Pseudomonas spp*., *Klebsiella spp*., *Salmonella spp*., *Bacillus spp. and Serratia spp*. based on biochemical and morphological characteristics. Of these six isolates, three isolates (EMBF2-*Klebsiella* spp, BCAF1- *Proteus* spp and BCAC2- *Bacillus* spp) were identify to solubilize phosphate by releasing a considerable amount of phosphate (12.01-21.23 ppm) and lowering the pH of the media. The three isolates showed tolerance to acidic and alkaline media and also showed plant growth promoting capabilities by releasing IAA and siderophores. The result revealed that the three isolates had potential to chelate the ion bond in Fe^3+^ in ferruginous ultisol by releasing low molecular weight organic acid, making phosphate to be bioavailable for plant usage. This will serve as biofertilizer in improving yield of crops in ferruginous ultisol and improve soil fertility.

## INTRODUCTION

Phosphorus is a major growth-limiting nutrient which plays important biochemical role in photosynthesis, energy storage and transfer, cell enlargement, cell division, N-fixation in legumes, respiration and several other processes in the living plant (Gyaneshwar *et al*., 2002). It has a concentration in the Earth’s crust of about one gram per kilogram. However, because of its reactive nature, phosphorous is not found free in nature, but is widely distributed in minerals, usually as phosphates (Mejia *et al*., 2016). Inorganic phosphate rock is today the chief commercial source of this element. However, it is not readily available for plant uptake unlike the case for Nitrogen (Ezawa *et al*., 2002). Phosphorus also have organic sources namely urine, bone ash and guano. These organic sources of phosphorus have been considered in previous researches as fertilizers in its pure form and also when mixed with compost (Adeoluwa *et al*., 2016). Soils are most times high in insoluble phosphate but deficient in bioavailable phosphate. Approximately 95–99% of soil phosphorous is present in the form of insoluble phosphates and hence cannot be utilized by the plants (Alok *et al*., 2013) especially in iron rich acidic soils such as ferruginous ultisols.

Ferruginous ultisol, also called red soils are iron rich acidic soils (Yu *et al*., 2016). This special soil landscape is primarily distributed throughout the tropical and subtropical areas, particularly in Southeast Asia, Oceania, South America, southern North America and Africa. (Zhao, 2014). The total area of red soils is approximately 64 million km^2^, accounted for 45.2% of the Earth’s surface area (Anumalla *et al*., 2019) and resided by 2.5 billion people, nearly half of the global population (Zhao, 2014). In Nigeria, it is predominant in some southern states such as Edo state, occupying about seven zones, including extreme north and central Benin (Dayou *et al*., 2017). Red soils are naturally poor in physical conditions and are also characterized by low pH, cation exchange capacity (CEC), fertility and difficulty to cultivate because of its low water holding capacity (Zhao, 2014). Red soils are rich in iron Fe^3+^ and low in bioavailable phosphate. This is because of the extreme reactive nature of phosphate anions (H_2_PO_4_^-^, HPO_4_^2-^) forming metal complexes with the rich iron (Fe^3+^) in ferruginous ultisols. These metal ion complexes precipitated about 80% of added P fertilizer and becomes unavailable to plants causing the anti-fertility nature of ferruginous ultisols (Goldstein, 1995).

Previously, farmers do apply phosphorus containing fertilizers in excess to increase the 20% of the bioavailable phosphorus for plant uptake (Yu *et al*., 2016), However, this process has led to chemical pollution and in turn decreased the soil fertility after long period of application and considering the high cost of the fertilizers, local farmers in some southern states of Nigeria such as Edo state, gave up on growing crops that are susceptible to phosphorus deficiency such as rice. Soil phosphorus dynamics involve mineralization and solubilization of inorganic phosphorus to bioavailable form by some heterotrophic microbes (Deubel *et al*., 2005). These microbes release low molecular weight metabolites such as gluconic and keto-gluconic acids (Goldsein, 1995: Deubel *et al*., 2005) which through their hydroxyl and carboxyl groups chelate the cation (Fe^+^) bound to phosphate and decrease the pH in basic soils (Stevenson, 2005). These heterotrophic microbes are the phosphate solubilizing bacteria (PSB). Due to the negative environmental impacts of chemical fertilizers and their increasing costs, the use of PSB is advantageous in the sustainable agricultural practices.

PSB can play an important role in supplementing phosphorus to the plants, allowing a sustainable use of phosphate fertilizers. Several strategies such as lowering of soil pH by acid production, ion chelation, and exchange reactions in the growth environment have been reported to play a role in phosphate solubilization by PSB (Alok *et al*. 2013). Considering the high ferruginous regions in Edo state and its low bioavailable phosphorus, PSB can be used to sustainably improve soil fertility and remediate iron toxicity. Also, a significant reduction in the use of phosphate fertilizer could be achieved if solubilization of soil-insoluble phosphorus is made available to crop plants (Thakuria *et al*., 2004). Hence, the current research aims at investigating the culturable bacterial as well as PSBs in ferruginous ultisol of Benin-city in comparative to control soils for future use as biofertilizer.

## MATERIALS AND METHODS

### Collection and analysis of soil samples

In the current study, six different rhizospheric ferruginous soils (F1-F6) were obtained at 0 - 20cm depth around Benin Metropolis. The control soil was collected from an area with high humus content around the rhizosphere of a mature banana plant. The physicochemical parameters of the soils were determined prior to usage of the soil according to standard procedures and presented in (Table 1). Soil pH was determined using a pH meter (Model PHS-3C) and soil organic matter was determined as percentage following Walkley and Black (1934). Soil available phosphorus was determined following Olsen method (Olsen *et al*., 1954). Available micronutrients such as Fe and Al were extracted from soil by DTPA according to Lindsay and Norvell (1978).

**Table 1:** Physicochemical properties of ferruginous and control soil used for the experiment

| Soil samples | pH | SOM (%) | Avl. P (mg/kg) | Fe (mg/kg) | Al (meq/100g) | WHC (%) |
| --- | --- | --- | --- | --- | --- | --- |
| F1 | 4.47 ± 0.50<sup>a</sup> | 1.50 ± 0.01<sup>a</sup> | 3.20 ± 0.01<sup>a</sup> | 200.12±3.09<sup>a</sup> | 15.02 ±0.50<sup>a</sup> | 26.93 ±0.02<sup>a</sup> |
| F2 | 6.81 ± 0.09<sup>b</sup> | 13.0 ± 0.05<sup>b</sup> | 22.85 ±0.20<sup>b</sup> | 72.13 ±0.35<sup>b</sup> | 2.02 ±1.41<sup>b</sup> | 48.44 ±0.01<sup>b</sup> |
| F3 | 5.62 ± 0.02<sup>c</sup> | 10.1 ± 0.64<sup>c</sup> | 11.43 ± 0.02<sup>c</sup> | 102.02 ±1.35<sup>c</sup> | 9.02 ± 0.50<sup>c</sup> | 48.43 ±0.01<sup>b</sup> |
| F4 | 5.82 ± 0.03<sup>d</sup> | 7.31 ± 0.09<sup>d</sup> | 13.21 ±0.05<sup>d</sup> | 90.10 ±0.40<sup>d</sup> | 6.23 ± 0.01<sup>d</sup> | 48.44 ±0.05<sup>b</sup> |
| F5 | 5.22 ± 0.02<sup>e</sup> | 5.99 ± 0.33<sup>e</sup> | 9.32 ± 0.02<sup>e</sup> | 112.30 ±0.05<sup>e</sup> | 10.71 ± 0.05<sup>c</sup> | 47.96 ±0.98<sup>b</sup> |
| F6 | 4.81 ± 0.01<sup>f</sup> | 2.30 ± 0.01<sup>a</sup> | 3.90 ± 0.03<sup>f</sup> | 190.20 ±4.46<sup>f</sup> | 12.55 ± 0.02<sup>c</sup> | 20.03 ± 0.01<sup>f</sup> |
| Control | 5.92 ± 0.98<sup>g</sup> | 17.08 ±0.09<sup>g</sup> | 20.21 ± 0.05<sup>g</sup> | 61.22 ±1.48<sup>g</sup> | 0.74 ± 0.05<sup>g</sup> | 85.11 ± 0.02<sup>g</sup> |
| Mean (F) | 5.08± 0.23 | 6.24± 0.92 | 9.91± 1.02 | 118.86± 12.44 | 8.61± 0.32 | 37.23± 4.02 |

SOM = soil organic matter, Avl. P = available phosphorus, Fe = iron, Al = aluminum, WHC=water holding capacity, F = ferruginous soil. Alphabet (a, b c) denotes significant difference when parameters for both soil samples are compared. Mean (F) = mean for each soil parameter obtained for ferruginous soil

### Microbial analysis

One (1) g of the collected soil samples were added to each of ten tubes containing 9ml distilled water thoroughly mixed and spread over petri plates containing Nutrient Agar, Actinomycetes, Isolation Agar such as MacConkey agar, Bacillus cereus agar, Pseudomonas cetrimide agar and Eosin methylene blue agar for enumeration of bacteria isolates. The plates were incubated at 30°C for 24 hours for bacterial isolation and at 30°C for 48 hours. Discrete colonies of viable bacteria cells on the medium were enumerated (counted) and the total heterotrophic bacteria count (THBC) was determined using the formula by Holtz (1993) as shown below:

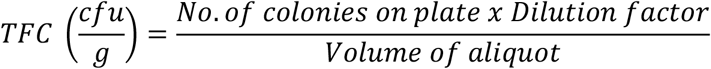

Soil microbial biomass carbon was measured through chloroform fumigation following Guang *et al*. (2012):

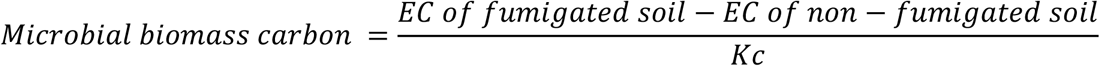

Where:

EC = Extractable organic carbon in fumigated soil and non-fumigated soil samples

Kc = 0.379, Kc is the K2SO4 extract efficiency factor for microbial carbon (Guang *et al*., 2012).

### Physiological Characteristics

The colony morphology of isolates was examined on nutrient agar plates. After an incubation of 3 days at 30°C, individual colonies were determined based on their colour on Mueller Hinton Agar (MHA), margin, elevation, shape, cell type, arrangement, spore staining, appearance, colony diameter, transparency and gram stain reaction (Aneja, 2003).

### Biochemical characteristics

Catalase test was performed by placing hydrolyzed H_2_O_2_ on a 2 days bacterial colony suspended with distilled water (Plate 1). Effervescence showed a positive result. Screening for ammonia production was performed by inoculation of freshly grown bacterial cultures in 10 ml nutrient broth and incubated at 34°C for 50 hours in a rotator shaker. After incubation, 0.5 ml of Nessler’s reagent was added to each tube. The development of a yellow to brown colour indicated a positive reaction for ammonia production following Holtz (1993). Nitrogen fixing activity was determined following Bashan and Holguin, (1998). Growth on nitrogen deficient medium confirms the ability to fix nitrogen following Bashan and Holguin (1998).

### Bromotyhmol blue test

This test was considered in order to determine the isolates with fast growth capacity. About 5 ml of 0.5% bromothymol blue solution was thoroughly mixed with 1L of nutrient agar. Bacteria isolates were streak on the plates and incubated at 30°C. The setup was monitored and recorded. Citrate utilization test was determined by inoculating fresh cultures on Simmons Citrate Agar slant and incubated for 24 hours at 370C. Appearance of blue colour indicated positive reaction, while a negative result obtained with no colour change. Oxidase test was performed using oxidase disc and observed for 15 seconds. Development of blue colour indicated a positive test. Other biochemical test such as sugar utilization activity, urease, lactose were determined following Kumar *et al*. (2014).

### Screening of Phosphate Solubilizing Bacteria (PSBs) from soil samples

Soil samples were subjected to serial dilution (10-folds) and then plated following the drop plate method on to a developed Pikovskaya’s growth medium as described by Pikovskaya (1948). These plates were subjected to aerobic incubation at 30°C for seven days. Appearance of holo-zone around the bacterial colonies was taken as indicator of phosphate solubilization (Plate 2). Colonies with obvious and clear zones were streaked again on Pikovskaya’s medium to obtain pure culture isolation. P-solubilization efficiency (PE) was also calculated (ppm) following Karpagam and Nagalakshmi (2014). While P-Solubilization index (SI) after 7 days was measured following:

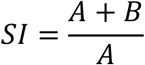

Where, A= Diameter of bacterial colony growth (mm)

B = Diameter of holo-zone (mm).

### Tolerance to different pH levels

This was determined by streaking bacterial isolates on Pikovskaya’s agar plates with pH adjusted to 4.0, 5.0, 6.0, 7.0, 8.0, 9.0 and 10.0 with HCl following (Mondala *et. al*., 2016).

### Screening for plant growth promoting (PGP) capabilities

The PGP capabilities of the PSB isolates were calculated by determining the IAA and Siderophores. The IAA was quantitatively determined using Luria Bertani broth following Gupta *et al*. (2012), while the siderophores production was quantitatively determined using MB medium and incubated for 72 hours following (Balkar, 2013).

### Statistical analysis

Data obtained were presented in means and standard errors. Data were analyzed following two-way analysis of variance using GENSTAT (8th edition). Where significant p-values were obtained, differences between means were separated using Student Newman Keuls test following (Alika, 2006).

## RESULTS AND DISCUSSION

### Soil physicochemical constituent of soil

Physicochemical properties of ferruginous and control soil used for the experiment have been presented on Table 1.

### Culturable bacterial composition of ferruginous ultisol, comparative to control soils

The mean total heterotrophic bacteria count (THBC) of the six ferruginous soils and the control soil sample used in the present study were presented in (Figure 1). The THBC ranged from 14.0 × 10^7^ to 0.20 × 10^7^ colony-forming units (cfu) per gram of soil. There were significant differences (P<0.05) in the average total bacterial counts of the different soil samples except for F1 and F4 soils which showed no significant difference (P>0.05). The highest bacterial count was observed in the control soil, followed by the F6 soil. However, the lowest bacterial count was observed in the F1 soil. This may be due to its acidic nature (4.47) and its other poor physicochemical conditions (Table 1) which alters bacterial growth. Microbial populations are influenced by environmental factors such as temperature and pH, moisture and organic matter (Fierer and Jakson, 2006). This result is consistent with the work of Smith *et al*. (2008) who suggested soil pH as the best predictor of bacterial community composition.

**Figure 1:**
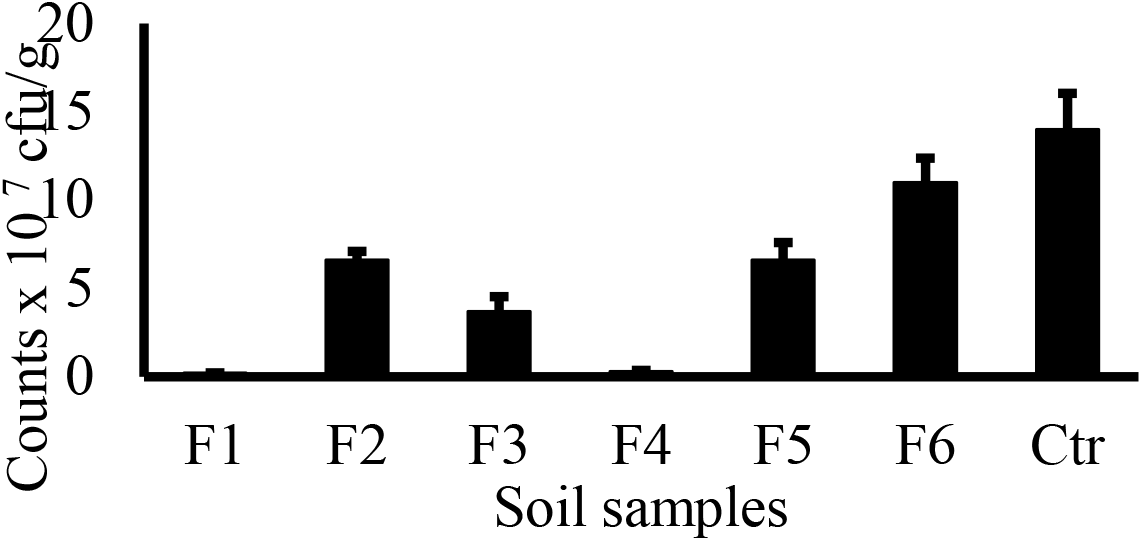
Average total heterotrophic bacterial count (THBC) of the sampling locations. F = Ferruginous soil, C= control soil.

### Soil microbial carbon

Soil microbial biomass carbon levels of the ferruginous soils and the control soils used in the present study was presented (Table 2). The control soil was observed to have the highest (380.01) soil microbial biomass carbon content, while the F1 soil had the lowest (10.40). There was significant difference in the soil microbial biomass carbon among all the sample sites except for F1 and F4. Since soil microbial biomass is a measure of the weight of microorganisms in soils (Musa, 2019), there is a significant positive correlation between the average soil THBC (Figure 1) and the microbial biomass carbon levels (Table 2).

**Table 2:** Microbial biomass carbon (MBC) levels

| Soil samples | MBC |
| --- | --- |
| F1 | 10.40 ± 0.41<sup>a</sup> |
| F2 | 103.20 ± 0.12<sup>b</sup> |
| F3 | 53.02 ± 1.2<sup>c</sup> |
| F4 | 12.32 ± 0.11<sup>a</sup> |
| F5 | 187.14 ± 0.39<sup>d</sup> |
| F6 | 210.20 ± 0.23<sup>e</sup> |
| Ctr | 380.01 ± 0.01<sup>f</sup> |
| p-value | 0.004 |

MBC = soil microbial biomass carbon, class, F = ferruginous soil, Ctr= control soil. Alhabet (a, b c) denotes significant difference when parameters for both soil samples are compared.

### Physiological characteristics of the isolates

The physiological characteristics of the isolates involving colony morphology was presented in (Table 3). The results indicated that all isolates have covex elevation, although some are having low convex. Only one isolate had smooth margin while the rest had entire margin. Also, all isolates showed cream colour on MHA and a circular shape. Gram staining test for all isolates showed negative except for one isolate (*Bacillus* spp.), while all isolates showed rod cell type. The microbial arrangement was single for all isolates except for one (*Proteus* spp.) which showed chains. Spore staining was not detected in all isolates except for one isolate (*Bacillus* spp.) which showed positive spore staining.

**Table 3:** Colony morphology of total heterotrophic bacteria

| **Cultural characteristics** |  |  |  |  |  |  |
| --- | --- | --- | --- | --- | --- | --- |
| Elevation | Low convex | Low convex | Convex | Convex | Convex | Convex |
| Margin | Smooth | Entire | Entire | Entire | Entire | Entire |
| Colour on MHA | Cream | Cream | Cream | Cream | Cream | Cream |
| Shape | Circular | Circular | Circular | Circular | Irregular | Circular |
| **Morphological** |  |  |  |  |  |  |
| Gram stain | Negative | Negative | Negative | Negative | Positive | Negative |
| Cell type | Rod | Rod | Rod | Rod | Rod | Rod |
| Arrangement | Chains | Single | Single | Single | Single | Single |
| Spore staining | ND | ND | ND | ND | Positive | ND |
| **Biochemical** |  |  |  |  |  |  |
| Catalase | + | + | + | + | + | + |
| Indole | + | - | + | - | - | - |
| Citrate | - | + | - | + | + | - |
| Urease | + | + | - | - | - | - |
| Oxidase | - | + | - | - | - | - |
| Glucose | + | - | + | + | + | + |
| Sucrose | + | - | + | + | + | + |
| Lactose | - | - | + | + | - | + |
| H<sub>2</sub>S | + | - | - | + | - | - |
| Gr. Diff. Agar | MCC (opaque) | PCA (green) | MCC (pink) | SSA (black) | BCA (Straw) | BCA (red) |
| Identity | *Proteus* | *Pseudomonas* | *Klebsiella* | *Salmonella* | *Bacillus* | *Serratia* |

Legend: MSA= Mannitol salt agar, PCA= *Pseudomonas* cetrimide agar; EMB = Eosin methylene blue agar, SSA = *Salmonella Shigella* agar, BCA= *Bacillus cereus* agar, MHA= Mueller Hinton Agar. ND=Not determined

### Biochemical characteristics of the isolates

All the distinct bacteria isolated showed positive to catalase test. However, all isolates showed different response to indole, citrate, urease, oxidase, glucose, sucrose, lactose and hydrogen sulphide. Furthermore, all isolates produced bubbles within 1 minute. This may be as a result of aerobic bacteria that hydrolyzes hydrogen peroxide, leading to the production of bubbles. Although, phosphate solubilizing bacteria may be aerobic or anaerobic. This result showed the bacteria isolated in this research are all aerobic bacteria.

Generally, a total of six distinct strains of bacteria (*Proteus* spp., *Pseudomonas* spp., *Klebsiella* spp., *Salmonella* spp., *Bacillus* spp. *and Serratia* spp.) were recorded from all the soil samples after accessing the cultural, morphological and biochemical characteristics. These bacteria have been isolated by previous studies from various soils in different agrogeology (Tani *et al*., 2011; Jahan *et al*., 2013; Swarnalakshmi *et al*., 2013; David *et al*., 2014; Min *et al*., 2016).

### Bromotyhmol blue test

Bacterial isolates from all ferruginous soil samples and the control developed a yellow colour (Plate 3). This indicates they are all acidic in nature (Table 4).

**Table 4:**
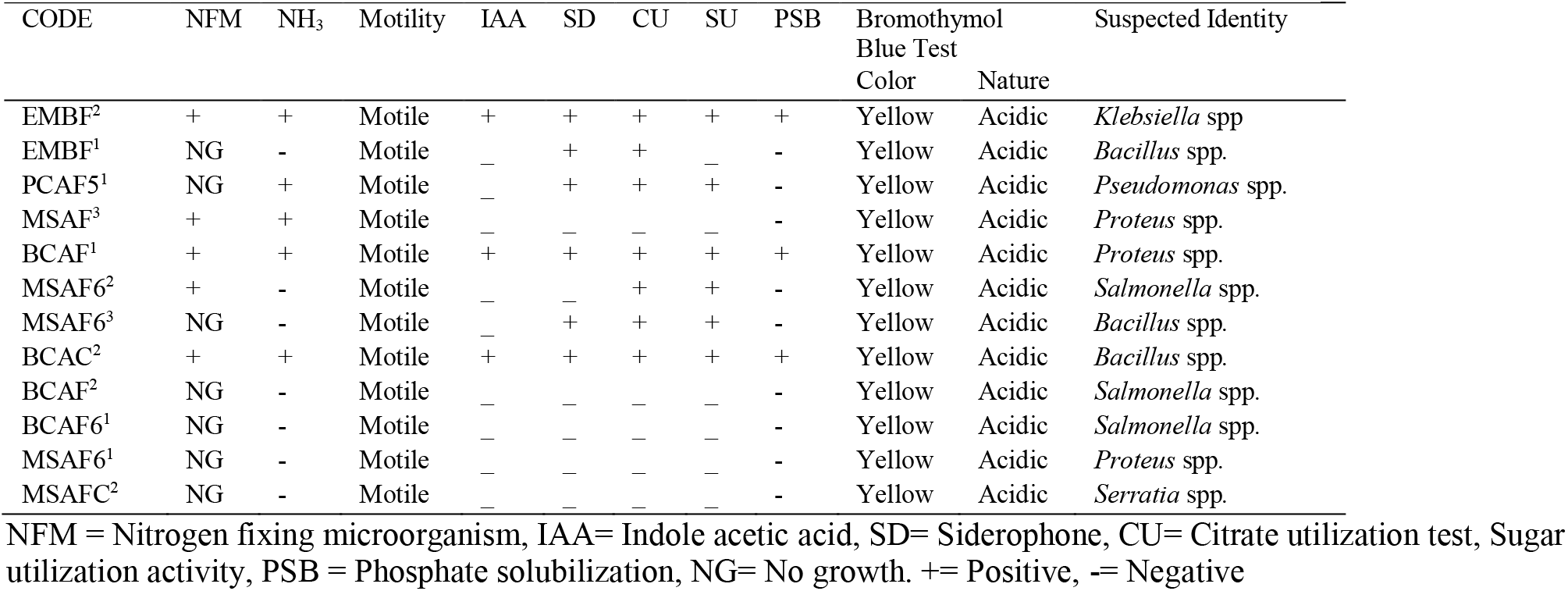
Physiological characteristic of isolated phosphate solubilizing bacteria

### Screening of Phosphate Solubilizing Bacteria (PSBs) from soil samples

The six distinct heterotrophic bacteria isolated from ferruginous soils and control soils were screened to determine their ability to solubilize phosphate based on halo-zone formation in Pikovskaya’s agar medium. Three (3) isolates proved positive to phosphate solubilization and were coded with (EMBF2, BCAF1 and BCAC2). These PSBs were obtained from F2, F1 and control soil respectively. Furthermore, the three isolates with PSB capability also showed plant growth promoting (PGP) characters such as Siderophores, indole acetic acid (IAA) and nitrogen fixation capabilities (Table 4). Many PSBs are reported as plant growth promoters (Musa, 2019; Hafeez, 2004). These isolates were selected for further analysis. Generally, isolates included: *Bacillus* spp., *Klebsiella* spp., *Proteus* spp., *Pseudomonas* spp., *Salmonella* spp., and *Serratia* spp.

**Plate 1:**
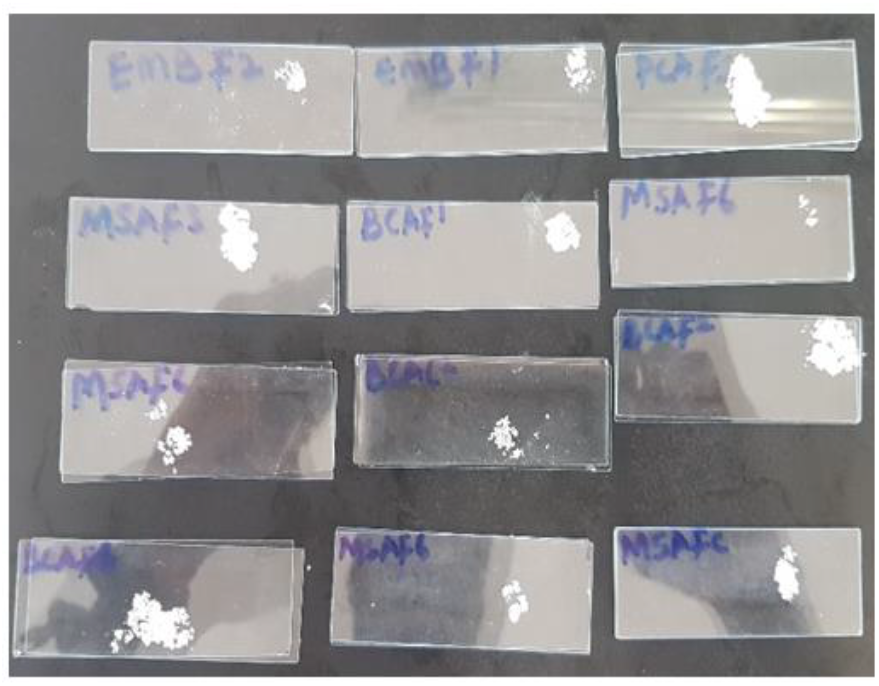
Qualitative catalase test

**Plate 2:**
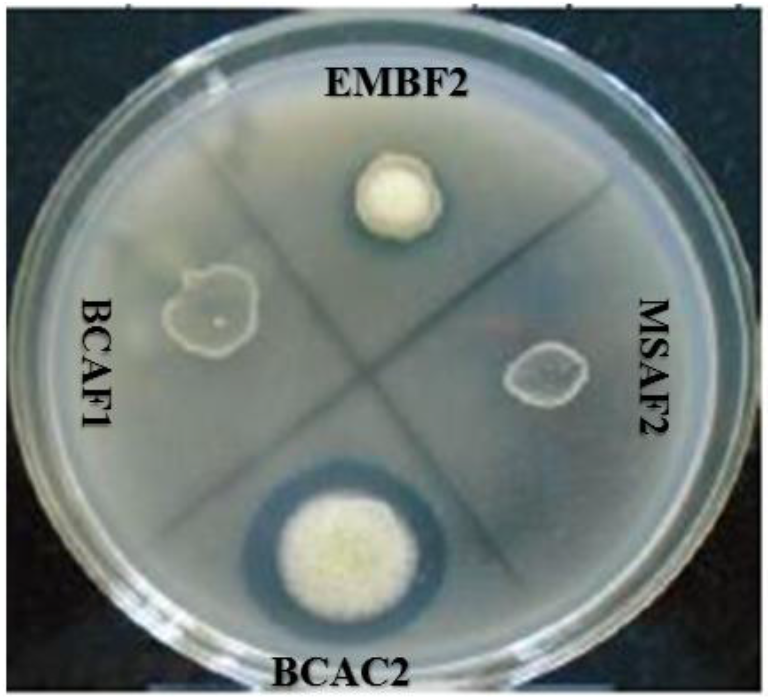
PSB forming clear holo-zone

**Plate 3:**
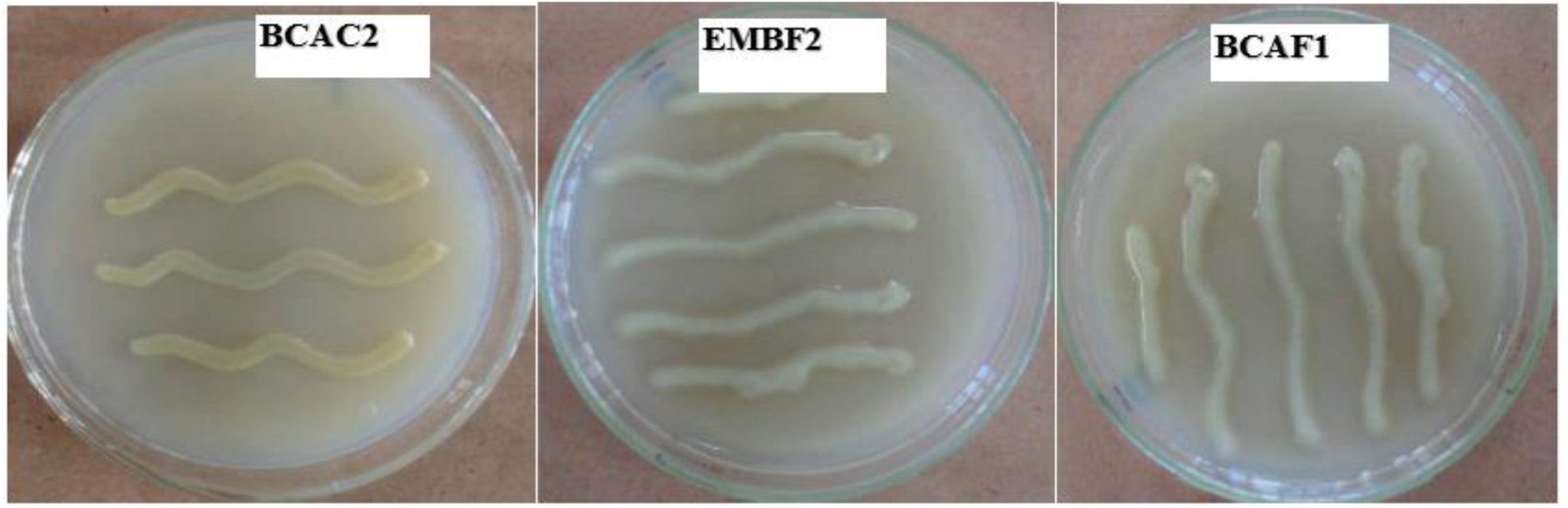
Bromothymol blue test of PSB isolates.

### PSB growth in different pH

PSB isolates were investigated for possible growth at different levels of pH in the Pikovskaya’s medium and presented in (Table 5). All PSB isolates grew in the medium with pH values of 5 to 8 although, the PSB from the control soil had higher growth at pH 4-6 while PSB from the F2 soils only had high growth at pH 6 compared to the PSBs from F1 soils. No PSB growth was detected in F2 and F1 soils at pH 9 and 10, indicating the isolates are not alkaline tolerant. However, there was PSB growth in the control soil at pH 9 but no growth at pH 10. This indicate that PSB in the control soil though tolerant to alkaline but its limited to pH 9 and can withstand acidic pH of up to 5. Furthermore, the result indicated that all the PSB isolates were tolerant to acidic nature, although PSB from the control soil showed stronger tolerance as against the PSB in the F1 and F2 soils. Soil pH values between 4 and 6 are best for P-availability, this is because these pH levels limits P from becoming fixed by iron in ferruginous soils, hence available for plant use. Azziz *et al*. (2012).

**Table 5:**
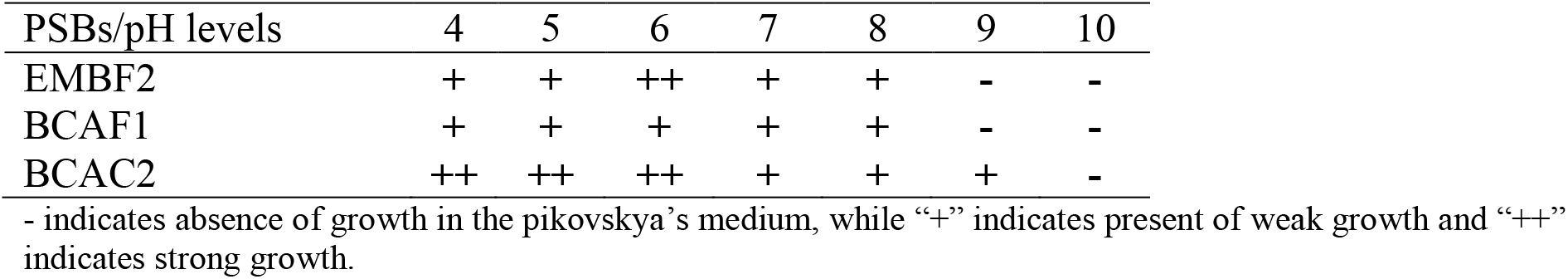
Tolerance of PSBs at different pH levels in Pikovskaya’s medium

### P-solubilization index

The PSB from the control soil showed highest holo-zone diameter (Plate 2) and bacteria colony diameter (32.4 ppm and 23.0 ppm respectively) on tricalcium phosphate. The BCAC2 isolate also showed the highest SI (2.4), followed by the EMBF2 isolate (2.34), while the BCAF1 had the lowest SI (2.17) in (Table 6). In this research, phosphate solubilization index is inversely proportional to pH levels (*R*^*2*^=0.9967) in (Figure 2). This result reaffirms that phosphate solubilization by PSB is involved in the production of organic acids (Halder *et al*., 1990; Rashid *et al*., 2004). This result is in accordance with the work of Fankem *et al*. (2006) who stated that phosphate solubilization is directly interpreted as the combined effect of pH decrease and organic acid production. (Table 5) has shown the capabilities of all isolates to withstand acidic pH, therefore the drop in pH as a result of solubilization process may not affect the isolates. Chen *et al*. (2006) reported that the solubilization of tricalcium phosphate in broth media by isolated strains of PSBs led to a significant decline in pH of the media from 6.8 to 4.9 after 72 hours as a result of the organic acid production by the PSB isolates which in turns, help in solubilizing phosphate.

**Table 6:** Solubilization of phosphorus by PSB isolates at pH 6.

| Isolates | Diameter of holo-zone (mm) | Diameter of bacterial colony (mm) | SI | pH |
| --- | --- | --- | --- | --- |
| EMBF2 | 23.6 | 18.4 | 2.28 | 4.1 |
| BCAF1 | 16.7 | 14.2 | 2.17 | 4.4 |
| BCAC2 | 32.4 | 23.0 | 2.40 | 3.7 |

SI= Solubilization index

**Figure 2:**
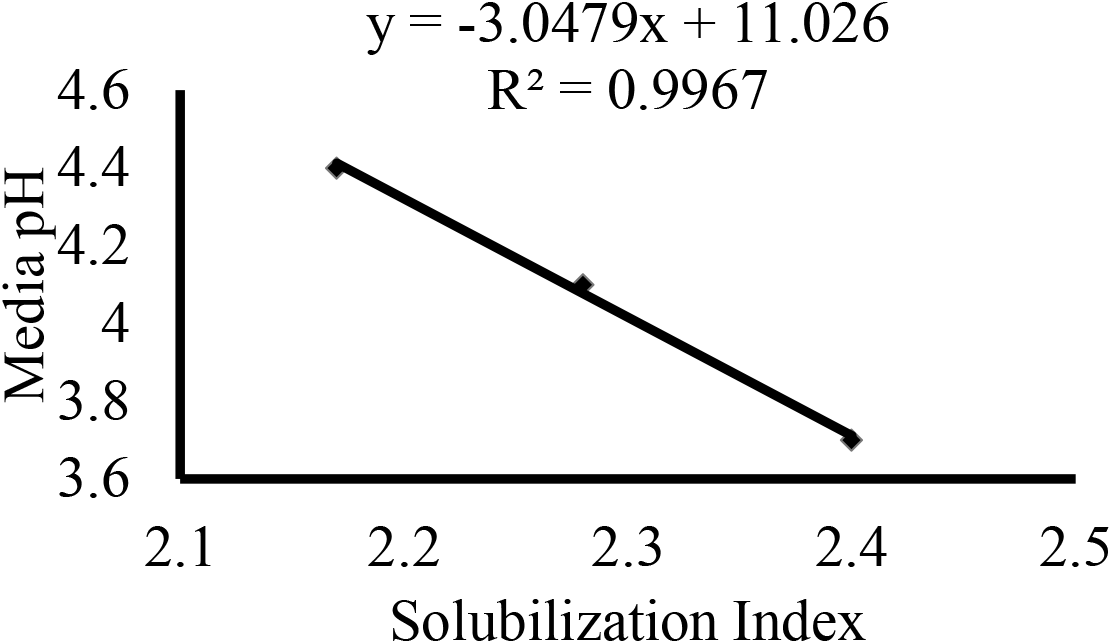
Relationship between solubilization index and pH of the Media.

P-solubilization efficiency of the three PSBs isolated from the ferruginous and control soils was presented in (Figure 3). This was obtained by growing the isolates on tricalcium phosphate containing media. The result revealed that phosphate solubilization efficiency varied between the three PSB isolated from the test soils. The PSB isolate from the control soil (BCAC2) showed the highest P-solubilization efficiency (21.23ppm), followed by the PSB from F2 soil (EMBF2-18.02ppm) while the PSB from F1 soil (BCAF1) had the lowest (12.01ppm) P-solubilization efficiency. Kundu *et al*. (2009) confirmed similar P-solubilization efficiency in the rhizosphere of chickpea.

**Figure 3:**
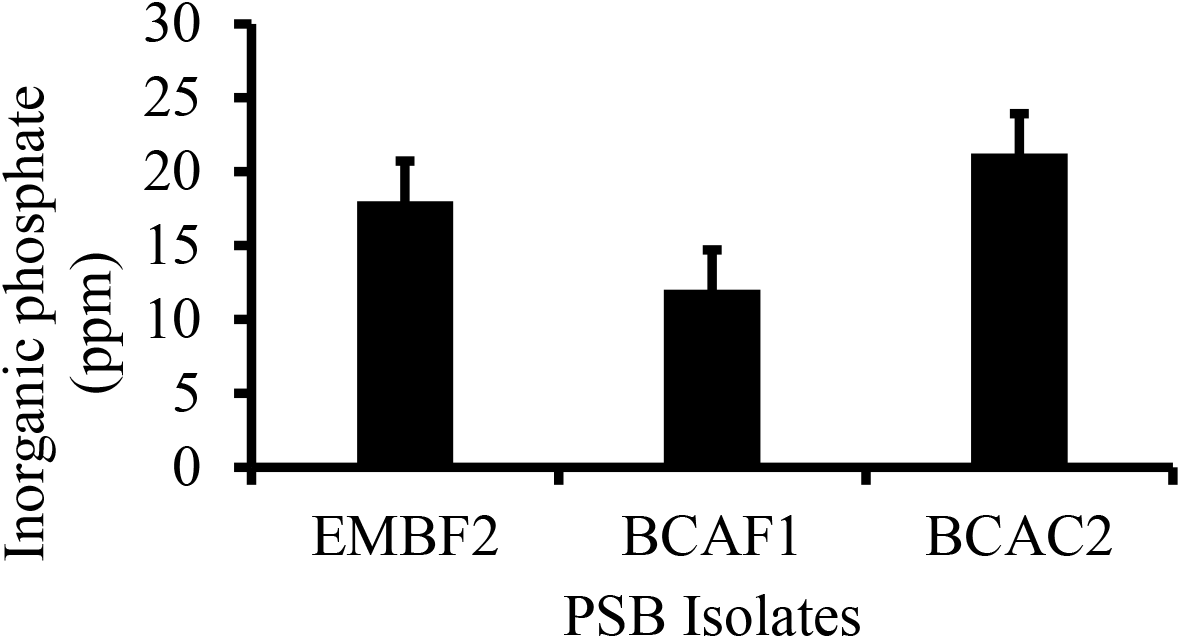
P-solubilization efficiency of PSB isolates

### Plant Growth Promoting (PGP) capabilities of the isolated phosphate solubilizing bacteria

Since many PSBs have PGP capabilities. The current study investigate the PSB capabilities of the isolated PSB by the production of IAA and siderophores. The result in Figure 4 and 5 showed that all the three isolates had the ability to produce IAA and siderophores. The siderophores was observed to range (74.4-20.4 %). BCAC2 was observed to produce the highest percentage siderophores, while the BCAF1 isolate had the lowest percentage siderophores. The quantitative result of IAA produced by the isolates showed the highest (29.4 µl/ml) was produced by the BCAC2 isolate, while the least IAA (21.0 µl/ml) was produced by the BCAF1 isolate. This result is consistent with work of Mamta *et al*. (2010); Beneduzi *et al*. (2013) and Wu *et al*. (2014) who used PSB as biofertilizers to improve the growth of *Stevia rebaudiana*, sugarcane and cotton respectively

**Figure 4:**
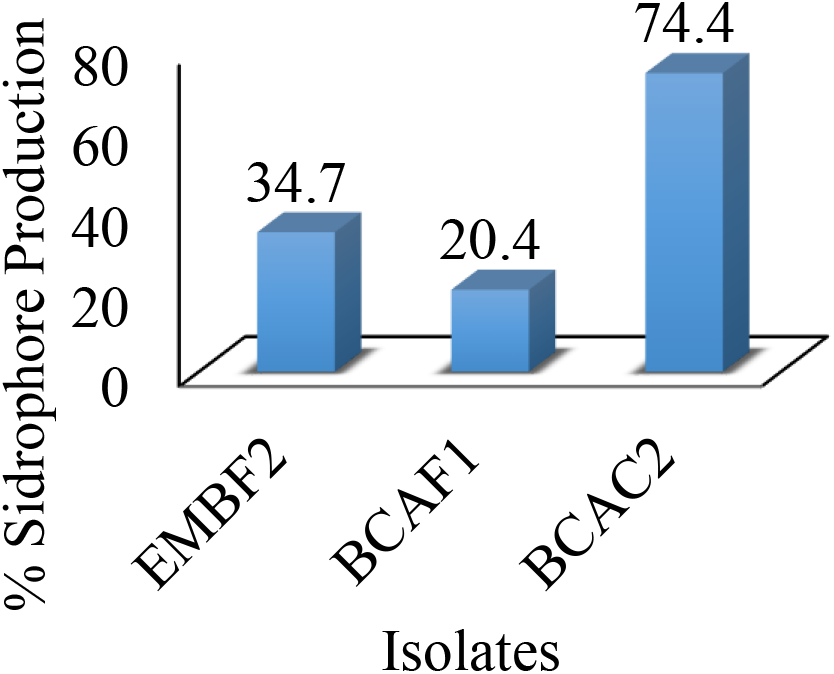
Percentage siderophores efficiency of the three isolated PSBs.

**Figure 5:**
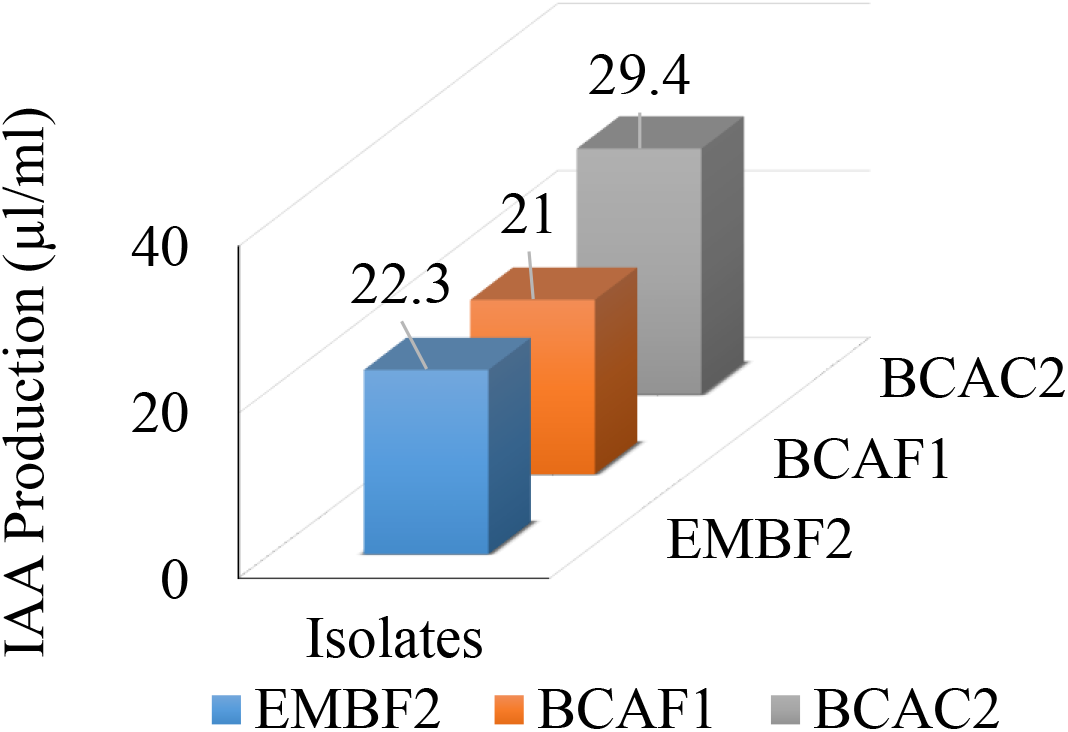
IAA production by the three isolated PSBs using Luria Bertani broth.

## Conclusion

Observations from the present research indicate that ferruginous soils are poor growing soils with less microbial composition compared to the control soil. Six distinct bacteria were isolated from the ferruginous and the control soils. A positive correlation was observed between the average soil THBC and the microbial biomass carbon levels. Three of these isolates (EMBF2-*Klebsiella* spp, BCAF1-*Proteus* spp and BCAC2-*Bacillus* spp) proved phosphate solubilization capabilities with the PSB from the control soil showing highest P-solubilization index. All the three PSB isolates showed tolerance to acidic and alkaline media of (4-9) pH. The P-solubilization efficiency of the isolates was proved by lowering the pH of the media at increasing SI. The Isolated PSB also showed PGP capabilities by releasing IAA and siderophores. Therefore, these isolates can be used to improve soil fertility by chelating the insoluble phosphorus and make it available for plant use in iron toxic soils such as ferruginous ultisol. This may also be employed as biofertilizers in order to improve growth of some agricultural crops in P-deficient and iron toxic soils. This will serve as a sustainable alternative to the inorganic p-fertilizers, reduce cost and improve yield of crops.

## Acknowledgements

The researchers are grateful to the Department of Plant Biology and Biotechnology, University of Benin, Nigeria and the Department of Biological Sciences, Admiralty University of Nigeria, Delta for the facilities. The mentorship and efforts of my supervisor, Beckley Ikhajiagbe, Ph.D., FIPMD, of the Department of Plant Biology and Biotechnology and the efforts of Dr. Abraham Ogofure of the Department of Microbiology, University of Benin, Nigeria during the course of the study is very much appreciated.

## Authors’ contributions

MSI and BI designed the study, MSI carried out the research under the supervision of BI. MSI carried out the statistical analysis and interpretation of data. MSI wrote the first draft. BI edited the final draft of the manuscripts. The authors read and approved the final manuscript.

## Funding

No funding was provided from any external source for the research. The research was sponsored by the authors.

## Consent for publication

Consent is given for publication of this manuscript when accepted.

## Conflict of interests

The authors declare no conflicts of interests.

## Notes

### Competing Interest Statement

The authors have declared no competing interest.

